# Development of an Intrinsic Skin Sensor for Blood Glucose Level with CRISPR-mediated Genome Editing in Epidermal Stem Cells

**DOI:** 10.1101/195529

**Authors:** Jiping Yue, Yuanyuan Li, Xuewen Gou, Xiaoyang Wu

## Abstract

Biointegrated sensors can address various challenges in medicine by transmitting a wide variety of biological signals. A tempting possibility that has not been explored before is whether we can take advantage of genome editing technology to transform a small portion of endogenous tissue into an intrinsic and long-lasting sensor of physiological signals. The human skin and the epidermal stem cells have several unique advantages, making them particularly suitable for genetic engineering and applications *in vivo*. In this report, we took advantage of a novel platform for manipulation and transplantation of epidermal stem cells, and presented the key evidence that genome-edited skin stem cells can be exploited for continuous monitoring of blood glucose level *in vivo.* Additionally, by advanced design of genome editing, we developed an autologous skin graft that can sense glucose level and deliver therapeutic proteins for diabetes treatment. Our results revealed the clinical potential for skin somatic gene therapy.

## Introduction

Biointegrated sensors can address various challenges in medicine by transmitting a wide variety of biological signals, including electrophysiological and biochemical signals continuously ^1^. Recent advancement in device designs and fabrication using novel nano-materials have greatly accelerated the development of new soft and stretchable electronics applicable *in vivo*, including brain, heart, and skin ^1,2^. However, it remains technically challenging in the field to improve the stability and biocompatibility of the device and resolve potential infection caused by device implantation. The recent development of genome editing technology ^3-5^ allows us to efficiently engineer cells and potentially transform a small portion of endogenous tissue into an intrinsic and long-lasting sensor of physiological and biochemical signals. However, this intriguing possibility has not been tested before.

The human skin and the skin epidermal stem cells ^6,7^ have several unique advantages, making them particularly suitable for genetic engineering and applications *in vivo* (Supplementary Fig. 1A). First, skin is the most accessible organ in our bodies, and its surface localization is ideal for signal transmission or monitoring. Secondly, the skin is one of the largest organs on the body surface, making it easy to isolate skin epidermal stem cells and to monitor the tissue for potential detrimental complications after engineering and, if necessary, to remove it in the case of an adverse consequence. The procedure to isolate and culture primary epidermal stem cells has been well established ^8-10^. Cultured epidermal stem cells can be induced to stratify and differentiate *in vitro*, and transplantation of epidermal autografts is minimally invasive, safe, and stable *in vivo.* Autologous skin grafts have been clinically used for the treatment of burn wounds for decades ^11,12^. However, despite of the potential clinical applications, the research of skin epidermal stem cells has been greatly hampered by the lack of an appropriate mouse model. Although it has been shown that mouse skin or human skin can be transplanted onto immunodeficient mice ^13-19^, the lack of an intact immune system in the host animals makes it impossible to determine the potential outcomes that the procedure may elicit *in vivo.* Immune clearance of engineered cells has been one of the major complications for somatic gene therapy ^20^. Additionally, it remains technically challenging to perform skin organoid culture with mouse epidermal stem cells and to generate mouse skin substitute for transplantation. To resolve these issues, we have developed a new organotypic culture model with mouse epidermal stem cells *in vitro* by culturing the cells on top of an acellularized mouse dermis ^21-23^. Exposing the cells to the air/liquid interface can induce stratification of the cultured cells to generate a skin-like organoid *in vitro* ^24^. Transplantation of the skin organoids to wild type (WT) mice with intact immune system has led to long-term and stable incorporation of engineered skin *in vivo* ^22^. In this report, with this unique isogenic skin transplantation platform, our results demonstrated that engineered skin grafts derived from genome-edited epidermal stem cells can serve as an internal biosensor for key biochemical signals, such as blood glucose, allowing long-term and non-invasive monitoring of glucose level *in vivo.*

## Results

### Glucose sensing with cultured mouse epidermal stem cells *in vitro*

Obesity and diabetes are acute and growing public health problem around the world ^25^. A biointegrated sensor for noninvasive monitoring of blood glucose level *in vivo* will remove the need for the patients to draw blood multiple times a day ^26-28^. Additionally, continuous monitoring of glucose allows the patients to better maintain the blood glucose level by altering the insulin dosage or diet according to the prevailing glucose values and prevent potential hyperglycemia and hypoglycemia. If the sensor can be connected with an insulin delivery device, it may create an “artificial endocrine pancreas” that can automatically maintain the glucose level in patients. Currently, most of the continuous monitoring sensors for glucose are enzyme electrodes or microdialysis probes implanted under skin ^26-28^. These sensors usually require oxygen for activity, and are insufficiently stable *in vivo* and poorly accurate under the low glucose condition. The presence of interfering electroactive substances in tissues can also cause impaired responses and signal drift *in vivo*, which necessitate frequent calibrations of current sensors. A fluorescence-based glucose sensor in the skin will likely be more stable, have improved sensitivity, and resolve the issue of electrochemical interference from the tissue ^29^.

To engineer epidermal stem cells for glucose sensing, we tested the intracellular expression of a sugar binding protein, glucose/galactose-binding protein (GGBP) ^30^. GGBP transports glucose within the periplasm of E.*coli*, and the binding of glucose to GGBP can lead to a large conformational change in the protein ^30^. This property of GGBP has been exploited to develop protein sensors for glucose using FRET (fluorescence resonance energy transfer) or bioluminescence imaging ^31-35^. WT (wild type) GGBP has a very high glucose binding affinity (*K*_*d*_ = 0.2 μm). To generate a probe corresponding to the physiologically relevant range of glucose, we engineered a CFP/YFP FRET sensor with A213R/L238S double mutant of *GGBP* (*K*_*d*_ = 10 mM) ^36^. Although skin stem cells are very susceptible for manipulation with viral vectors, viral infection could lead to genotoxicity and may raise a significant safety concern for the potential applications *in vivo* ^37,38^. The recent advancement of genome editing technology with CRISPR system presents a novel approach to carry out a site-specific modification of the target genome non-virally ^5,39,40^. To test CRISPR-mediated genome editing in mouse epidermal stem cells, we developed DNA vectors encoding the D10A mutant of *Cas9* (CRISPR associated protein 9) ^41^, two gRNAs (guide RNA) targeting the mouse *Rosa26* locus, and a *Rosa26*-targeting vector. The targeting vector contains two homology arms for the *Rosa26* locus, flanking an expression cassette that encodes a *GGBP* fusion protein (Fig. 1A and Supplementary Fig. 1B).

**Figure 1.**
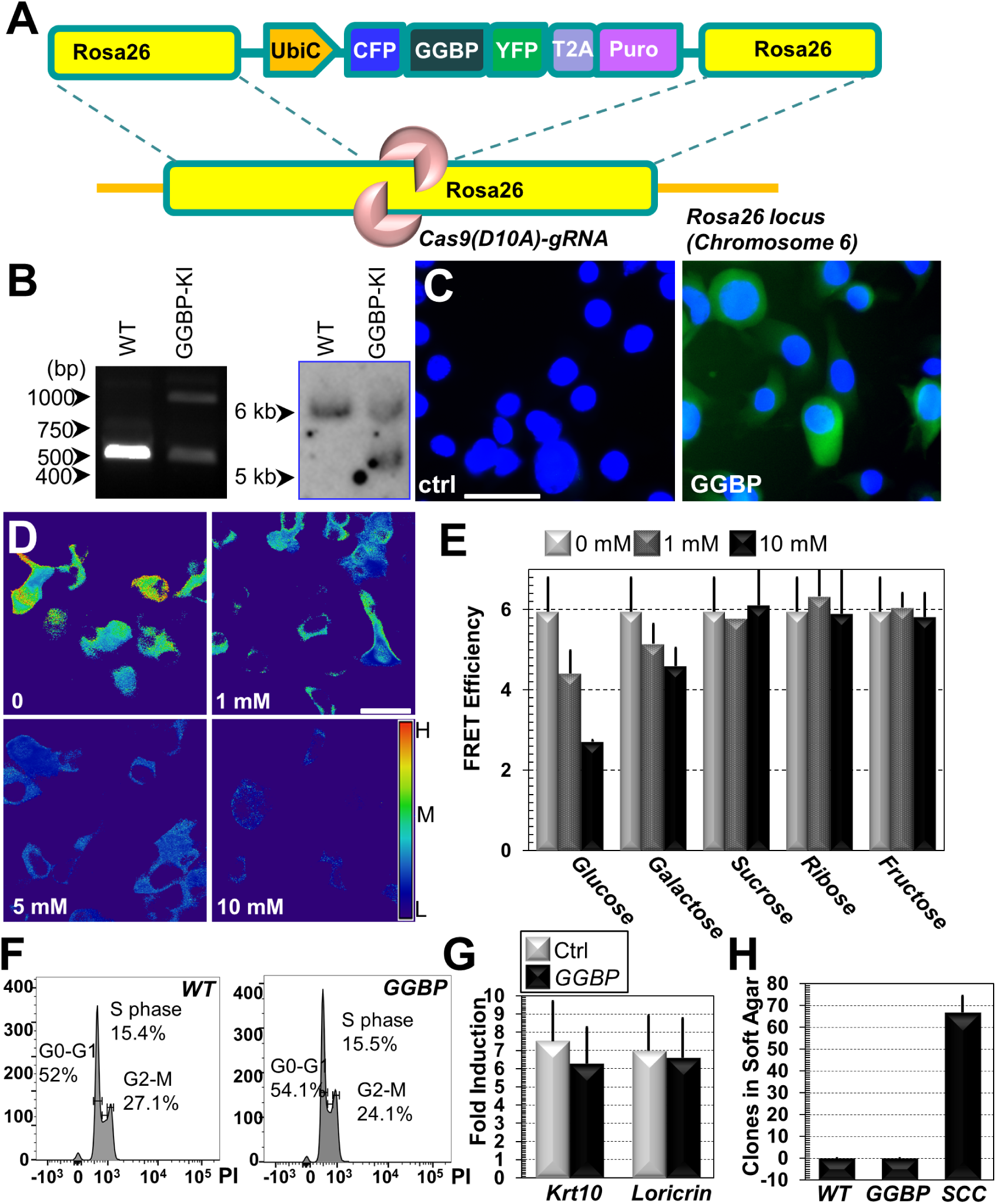
Engineering skin epidermal stem cells with CRISPR. **(A)** Diagram showing the *Rosa26* targeting strategy for the expression of glucose sensor *GGBP.* The targeting vector contains two *Rosa26* homology arms, flanking the expression cassette for *GGBP* and a selection marker (puromycin resistant gene, Puro) driven by a constitutive promoter UbiC (Ubiquitin C promoter). GGBP and Puro are separated by a self-cleavable peptide *T2A.* **(B)** Integration of the targeting vector into *Rosa26* locus was verified by PCR (left panel) and southern blot (right panel). Positive clones displayed an additional band of the expected size. **(C)** Expression of the *GGBP* sensor in the targeted cells was confirmed by fluorescence imaging. Scale bar=20μm. **(D)** FRET ratio images are pseudocolored to demonstrate glucose-dependent ratio changes in the engineered cells. Red indicates high (H) FRET efficiency, and blue represents low (L) efficiency. M: medium FRET efficiency. Integration of the ratio over the entire cells was used to quantify the FRET ratio changes. **(E)** The FRET ratio change of the GGBP reporter was determined in the presence of various monosaccharides or oligosaccharides at different concentration. Note only glucose and galactose lead to significant FRET ratio changes (*P* < 0.01). *n* > 6 (individual cells). **(F)** FACS (fluorescence activated cell sorting) demonstrated similar cell cycle profiles in WT (wild type) and *GGBP-*expressing epidermal stem cells. PI: propidium iodine. **(G)** Western blot analysis of early (Krt10) and late (loricrin) differentiation marker expressions in WT and *GGBP-*expressing cells upon calcium shift. Band intensity was determined by densitometry and the fold of induction is quantified. *n* = 4 (4 independent tests). *P >* 0.05. **(H)** WT cells or *GGBP* cells were tested for anchorage independent growth in soft agar. Note no growth for WT or GGBP cells, but tumor initiating cells isolated from skin SCC (squamous cell carcinoma) can readily produce colonies in soft agar plate. *n* = 3. *P <* 0.01 (between WT and SCC or GGBP and SCC).

Primary epidermal keratinocytes were isolated from CD1 newborn mice, and were electroporated with the *Rosa26* targeting vector together with the plasmids encoding *Cas9* and *Rosa26-*specific gRNAs. Clones were isolated upon selection and the correct integration into the *Rosa26* locus was confirmed by both PCR screening and southern blot analysis (Fig. 1B). Engineered epidermal cells exhibited robust expression of *GGBP* fusion proteins in the cytosol (Fig. 1C). To test the glucose sensing *in vitro*, we exposed the cells to medium containing an increasing amount of glucose. Quantification of FRET ratio by microscopic imaging showed an excellent correlation of FRET ratio with extracellular glucose concentration, ranging from 0-10 mM (Fig. 1D). Our analysis further showed that the GGBP reporter responded to extracellular glucose or galactose, but not to other sugars, such as sucrose, fructose, or ribose (Fig. 1E). The intracellular GGBP probe responded to the change of glucose concentration rapidly. The FRET ratio changed within 30 seconds after replacement of the medium, and remained stable in the same medium. To assess the intracellular GGBP probe more quantitatively, the reporter-expressing cells were perfused with an increasing concentration of glucose. When a stable FRET ratio was reached for each concentration, glucose was removed by perfusion and exchanged with a glucose-free medium. Perfusion of glucose from 0.1 mM to 10 mM led to reversible and increasing FRET efficiency changes (Supplementary Fig. 1C). When exposed to glucose at higher concentration, the FRET efficiency changes followed a hyperbolic curve, reflecting saturation of the probe at concentration higher than 25 mM (Supplementary Fig. 1D). Together, our results suggest that epidermal stem cells expressing the *GGBP* reporter can faithfully sense the extracellular glucose concentration.

Expression of the *GGBP* fusion protein in epidermal cells did not significantly affect cell proliferation (Fig. 1F and Supplementary Fig. 1E) or differentiation (Fig. 1G) *in vitro.* To confirm that modified epidermal cells are not tumorigenic, we examined the potential anchorage-independent growth of the modified cells. Our results indicate that epidermal stem cells with the *GGBP* reporter cannot grow in suspension (Fig. 1H). As a positive control, cancer initiating cells ^42^ isolated from the mouse SCC (squamous cell carcinoma) exhibited robust colony formation in soft agar medium (Fig. 1H). Expression of the *GGBP* reporter did not affect the ability of epidermal stem cells to stratify. When subjected to skin organoid culture ^22^, the targeted cells readily produced stratified epithelial tissue (Supplementary Fig. 1F).

### Glucose sensing with skin transplants *in vivo*

To investigate the potential applicability *in vivo*, we prepared the skin organoid culture with the engineered epidermal stem cells ^22^, and transplanted the organoids to CD1 host animals (Fig. 2A). No significant rejection of the skin grafts has been observed after transplantation, suggesting that the targeted epidermal stem cells are well tolerated immunologically *in vivo.* The grafted skin exhibited normal epidermal stratification, proliferation and cell death (Supplementary Fig. 2A-E). To test the glucose sensing capability *in vivo*, we carried out the IPGTT (intraperitoneal glucose tolerance test) in grafted animals. Fasted animals were administered with a bolus of glucose intraperitoneally. Fluorescence (FRET) change in the grafted skin was monitored by intravital imaging (Fig. 2B), and the blood glucose level was measured by a commercial glucose monitoring system (Bayer Contour) with the blood samples taken from the snipped tail. Figure 2C and D showed the correlation between the measured glucose concentration and the FRET ratio changes over time. Interestingly, the FRET ratio exhibited a nearly linear correlation with the glucose concentration *in vivo* (Fig. 2D, R^2^=0.977).

**Figure 2.**
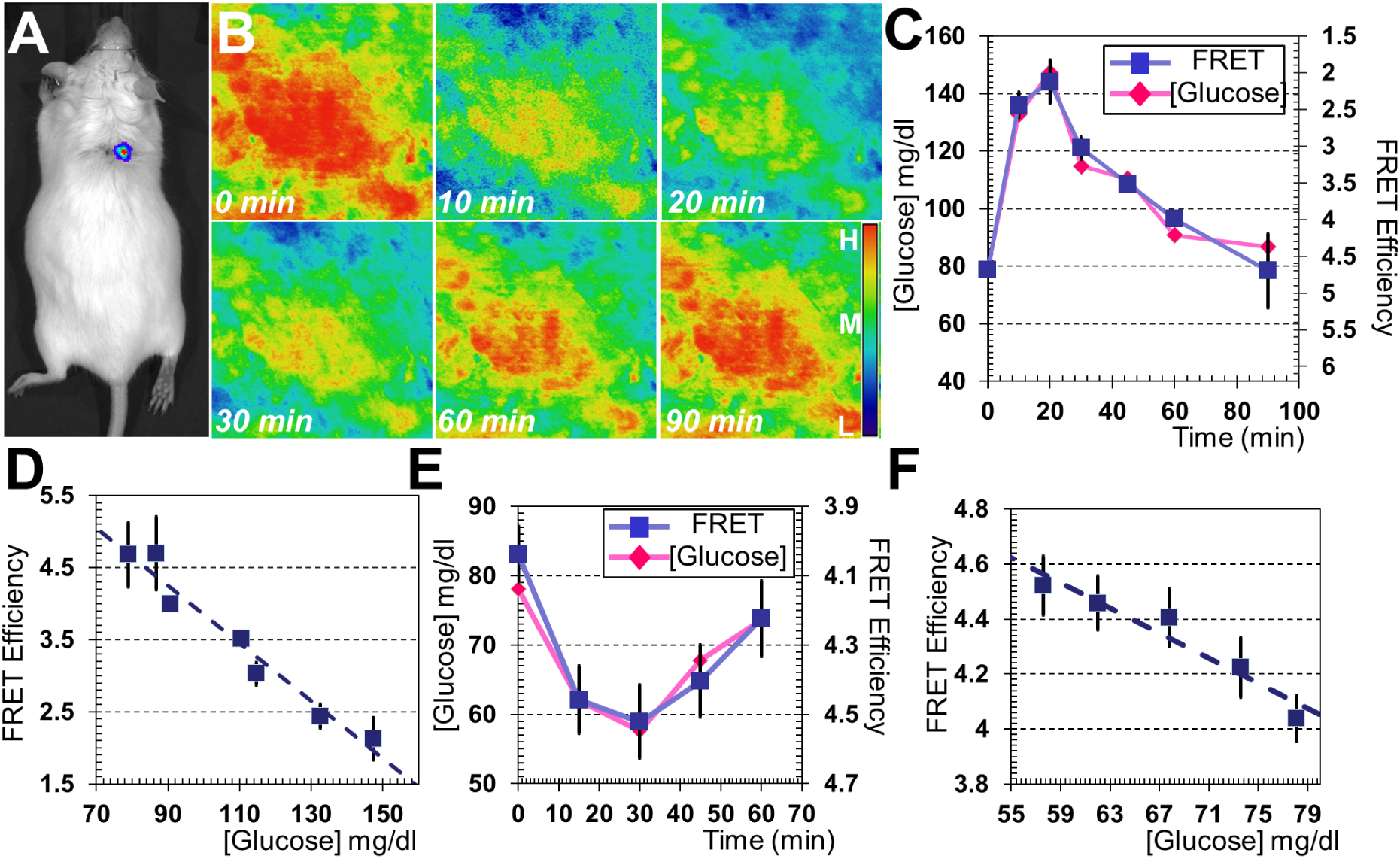
Monitoring changes of blood glucose level with GGBP reporter *in vivo*. **(A)** Skin organoids were developed from control or *GGBP*-producing cells, and were transplanted to CD1 mice. **(B)**Glucose fluctuation was induced in the grafted animal with IPGTT (intraperitoneal glucose tolerance test). FRET ratio images were pseudocolored to demonstrate glucose-dependent ratio changes in the grafted skin. Red indicates high FRET efficiency, and blue represents low efficiency. **(C-D)** Correlation of FRET ratio with blood glucose concentration upon IPGTT (intraperitoneal glucose tolerance test). *n* = 9 (integrated FRET value from 9 different fields at each time point). **(E-F)** Insulin injection was used to induce hypoglycemia in the grafted animals. Intravital imaging of the grafted skin demonstrated the correlation between FRET ratio of the GGBP reporter with the blood glucose concentration. *n* = 9 (integrated FRET value from 9 different fields at each time point).

Traditional glucose sensor cannot accurately measure low glucose level *in vivo* ^26,27^. To test the skin sensor under lower glucose conditions, we induced hypoglycemia by insulin administration to fasted animals. Intravital imaging of the grafted skin again showed excellent correlation of the FRET ratio changes with glucose levels (Fig. 2E and F). To further evaluate the utility of our sensor in a physiological setting, fasted mice were fed with standard chow diet for 30 minutes, and blood glucose level was monitored afterwards. Our results demonstrated the correlation between the blood glucose levels with FRET efficiency changes (Supplementary Fig. 2F-G). The reporter also showed excellent dynamic range when hyperglycemia was induced *in vivo* by intraperitoneal injection of higher dose of glucose (Supplementary Fig. 2H). Together, our results strongly suggest that the engineered skin graft with the GGBP reporter can accurately sense the blood glucose level *in vivo.*

### Glucose sensing and delivery of therapeutic proteins with a single skin transplant

The unique advantage of our epidermal stem cell platform allows complex genetic engineering to be applied with ease. Thus, introduction of an expression cassette that encodes both *GGBP* reporter and a therapeutic protein into the epidermal stem cells may achieve continuous glucose monitoring and diabetes treatment with one single skin transplantation. The hormone GLP1 (glucagon like peptide 1) is released from the gut upon food intake. GLP1 acts both as a satiety signal to reduce food consumption and as an incretin hormone to stimulate insulin release and inhibit glucagon secretion ^43^. These benefits have made GLP1 receptor agonists being used to treat type 2 diabetes. Our recent study demonstrated that skin-derived GLP1 release can effectively reduce body weight and inhibit diabetes development in a high fat diet model ^22^. To develop skin transplant with multiple functionalities, we engineered a new *Rosa26* targeting vector containing an expression cassette that encodes a *GLP1* and mouse *IgG-Fc* fragment fusion protein together with the *GGBP* reporter (Supplementary Fig. 3A). Fusion with IgG-Fc can enhance the stability and secretion of GLP1 when ectopically expressed in cells ^44^. The engineered epidermal cells exhibited a robust GLP1 production and secretion *in vitro* and *in vivo* (Fig. 3A and Supplementary Fig. 3B). Since the ectopic expression of *GLP1* is driven by a constitutive promoter, the expression and secretion of *GLP1-Fc* fusion protein by the skin epidermal cells led to a constitutively elevated level of GLP1 in blood regardless of nutrient or food intake (Supplementary Fig. 3C). The secreted GLP1 fusion protein is functional as the conditioned medium from *GLP1-*expressing cells can significantly induce secretion of insulin when added to insulinoma cells cultured *in vitro* (Fig. 3B).

**Figure 3.**
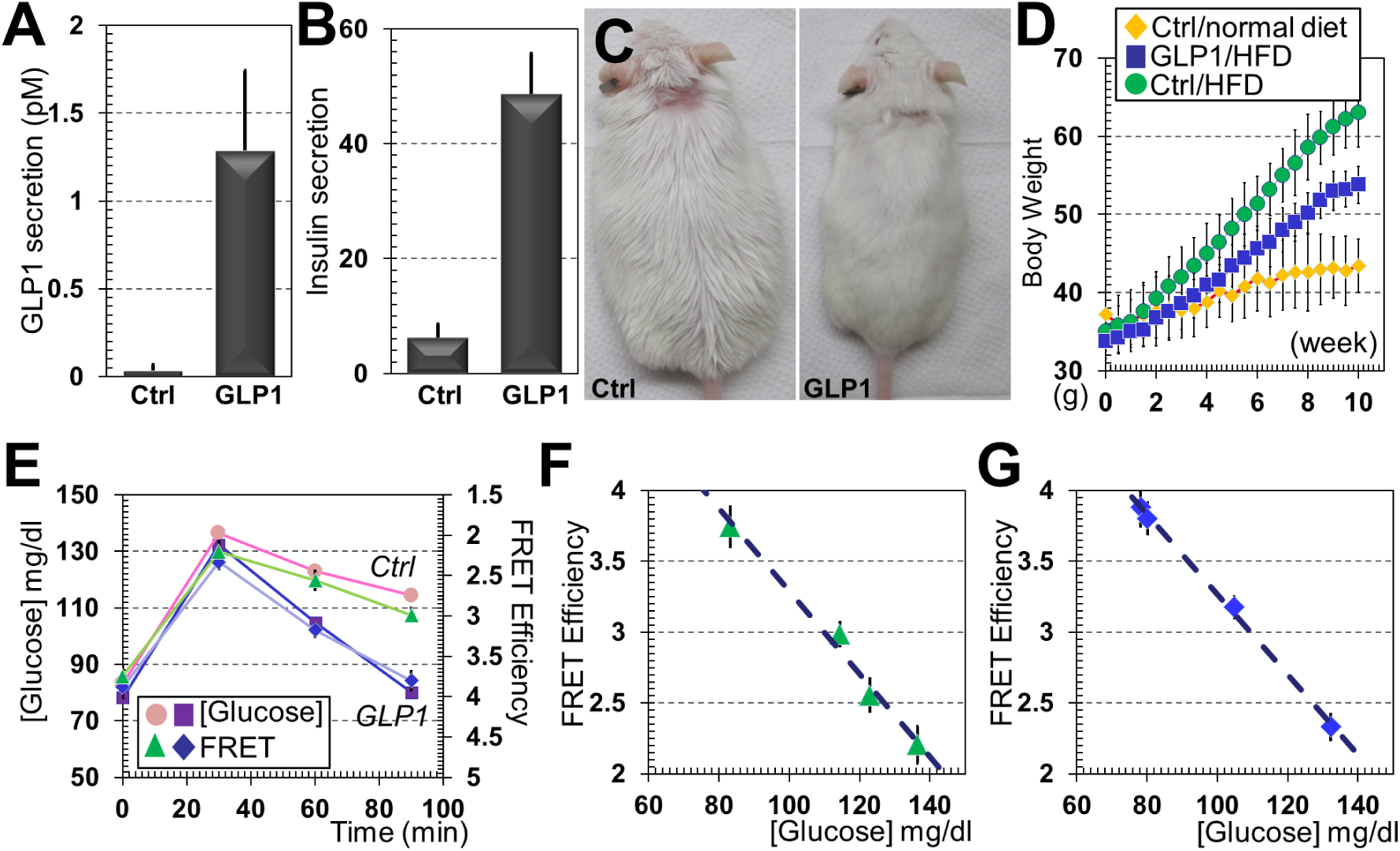
Monitoring blood glucose level and delivery of therapeutic molecules with single skin transplant. **(A)** Secretion of *GLP1* in cell culture medium was determined by ELISA (enzyme-linked immunosorbent assay). *n* = 3 (3 independent tests). *P* < 0.01. **(B)** Conditioned medium was collected from different cell cultures and used to treat starved insulinoma cells. Secretion of insulin *in vitro* was determined by ELISA. *n* = 4 (4 independent tests). *P* < 0.01. **(C)** Images of control and GLP1 animals fed with HFD (high fat diet). **(D)** Body weight change of different cohorts of mice measured from ∼10 weeks of age. Note that HFD induced significant obesity in control mice (*P* < 0.01, between control mice with normal diet or HFD for week 8-10) but expression of *GLP1* inhibited weight gain (*P* < 0.05, between GLP1/HFD and control/HFD groups for week 8-10). *n* = 5 (animals). **(E)** Correlation of FRET ratio with the blood glucose concentration over time upon IPGTT in control and GLP1 mice. *n* = 9 (integrated FRET value from 9 different fields at each time point). **(F-G)** Correlation of blood glucose concentration with GGBP FRET changes in control (L) and GLP1 (M) mice. *n* = 9 (integrated FRET value from 9 different fields at each time point).

To examine the potential applicability in diet-induced obesity and diabetes, we transplanted *GLP1/GGBP-*expressing cells and control cells (*GGBP* alone) to two cohorts of CD1 adult mice, and employed high fat diet (HFD) to induce obesity in grafted animals. To minimize gender difference, only male animals were used in our study. Compared with the animals on regular chow diet, the HFD greatly accelerated body weight gain in the mice grafted with control cells. By contrast, *GLP1* expression led to a significant inhibition in body weight increase (Fig. 3C and quantification in 3D). To examine glucose homeostasis, we conducted IPGTT. Expression of *GLP1* significantly reduced glycemic excursion *in vivo* as determined by both direct measurement of blood glucose or intravital imaging of the *GGBP* reporter (Fig. 3E-G). Noninvasive monitoring of the GGBP reporter exhibited an excellent correlation with the conventional glucose measurement in both diabetic animals and the GLP1-treated animals (Fig. 3F and G).

### Glucose sensing with human epidermal stem cells

To test the feasibility of glucose sensing with human epidermal stem cells, we cultured human skin organoids from primary epidermal keratinocytes isolated from human newborn foreskin. The human epidermal keratinocytes can readily produce organoids *in vitro,* which can be transplanted to the *nude* mice. When infected with lentivirus, the grafted human cells exhibited robust expression of the exogenous *Luciferase* gene (Fig. 4A). The grafted tissue showed normal skin stratification when stained for early or late epidermal differentiation markers (Fig. 4B). To examine CRISPR-mediated genome editing in human epidermal cells, we developed vectors encoding two gRNAs targeting human *AAVS1* (adeno-associated virus integration site 1) locus, and an *AAVS1*-targeting vector (Supplementary Fig. 4A) that encodes *GGBP* reporter protein. Human epidermal keratinocytes were electroporated with the targeting vector together with the plasmids encoding *Cas9* and the gRNAs. Clones were isolated and the correct integration was confirmed by southern blot analysis (Fig. 4C). Expression of the *GGBP* fusion protein in human cells did not significantly affect cell proliferation (Supplementary Fig. 4B) or differentiation (Supplementary Fig. 4C) *in vitro.* The engineered cells stratified and formed skin organoids *in vitro,* which were transplanted to the *nude* host. The grafted skin exhibited normal epidermal stratification and proliferation *in vivo* (Supplementary Fig. 4D-G). With IPGTT in the grafted nude mice, we carried out intravital imaging of the GGBP reporter (Fig. 4D) and observed similar correlation of FRET ratio changes in the grafted skin with blood glucose levels (Fig. 4E and F). Our data strongly suggest that the skin-based glucose sensor can be used to monitor blood glucose level in human patients.

**Figure 4.**
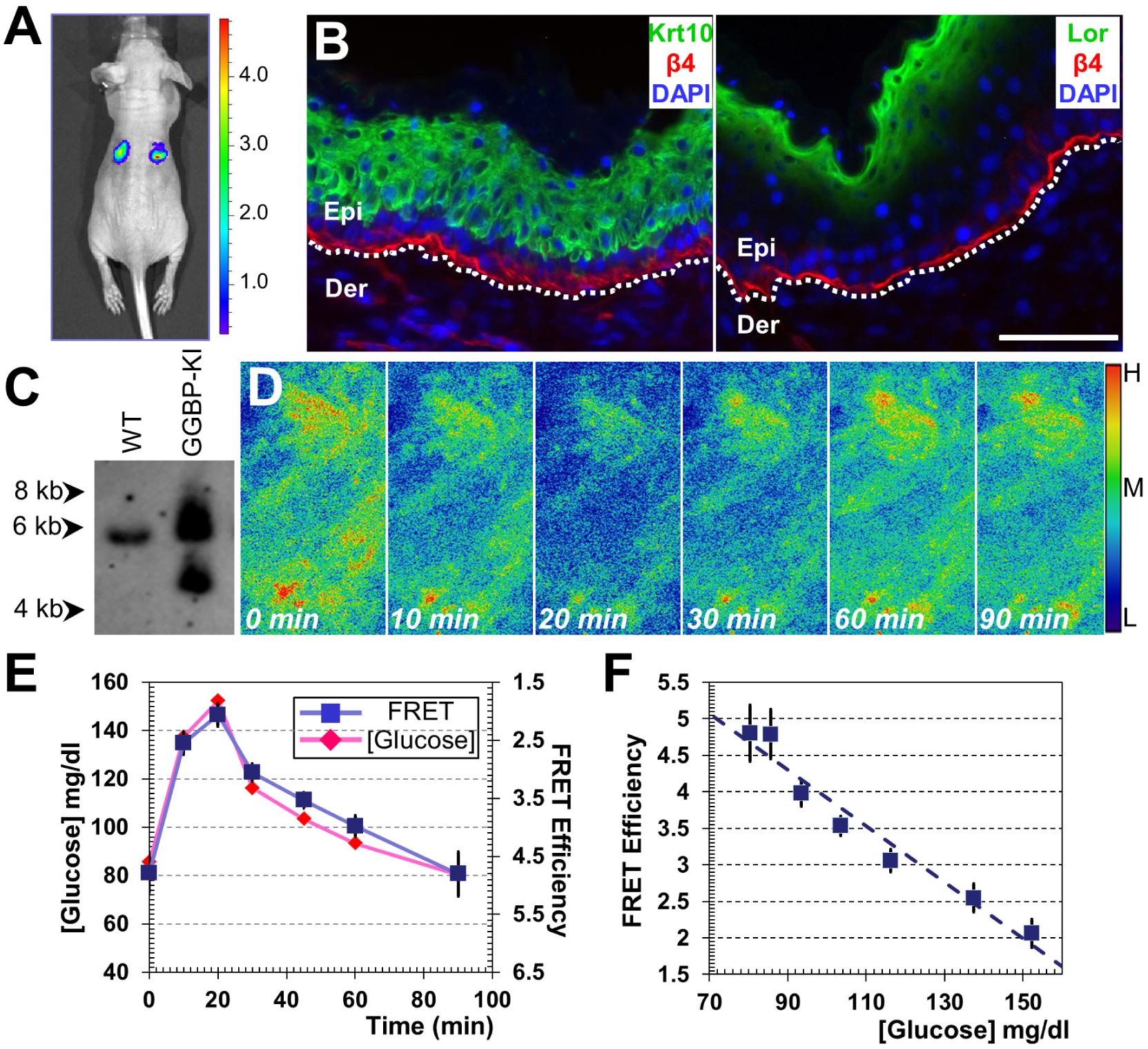
Expression of *GGBP* reporter in human epidermal stem cells with CRISPR. **(A)** Image of nude mouse grafted with organotypic human skin culture. Intravital imaging showed efficient incorporation of grafted cells that express *luciferase*. **(B)** Sections of grafted skin and the adjacent host skin were immunostained with different antibodies as indicated. Scale bar=50 μm. **(C)** Integration of the targeting vector into *AAVS1* locus was verified by southern blot. Positive clones displayed an additional band of the expected size. **(D)** Glucose fluctuation was induced in grafted animal with IPGTT (intraperitoneal glucose tolerance test). FRET ratio images were pseudocolored to demonstrate glucose-dependent ratio changes in the grafted skin. Red indicates high FRET efficiency, and blue represents low efficiency. **(E-F)** Correlation of FRET ratio with the blood glucose concentration upon IPGTT. *n* = 9 (integrated FRET value from 9 different fields at each time point).

## Discussion

Noninvasive monitoring of blood glucose level in a continuous manner can provide significant clinical benefits to the diabetic patients ^26-29^. The current methodologies of continuous glucose detection mostly depend on electrochemical approaches (enzymatic or non-enzymatic approaches) and optical approach (fluorescence, FRET, ocular spectroscopy, infrared spectroscopy, photoacoustic spectroscopy, *et al.*) ^26,28,45-47^. In order to sense the glucose level *in vivo,* most current technologies have to use invasive sensors that are implanted subcutaneously or intravenously. However, implantation of the devices can lead to significant tissue inflammation and biofouling-induced sensor degradation, causing loss of sensitivity. Although less invasive approaches have been developed, such as iontophoresis, sonophoresis, smart tattoo, and microneedles, these technologies suffer from various drawbacks, including calibration inaccuracy, biological incompatibility, current-induced skin erythema or irritation, significant subject-to-subject variability, and long warm-up or duration time required to collect sufficient amount of sample ^26,28,29,45-48^. Thus, the development of novel, noninvasive, reliable, and long-lasting sensor for continuous glucose monitoring will make a tremendous contribution to the field.

Because epidermal stem cells allow efficient genetic manipulation with the minimal risk of tumorigenesis or other detrimental complications *in vivo* ^13,14^, the skin may serve as an ideal intrinsic sensor of our own biochemical and physio-pathological signals. We have recently resolved the technical hurdles and established the unique mouse-to-mouse skin transplantation model with immunocompetent hosts ^22^. Our results provide the key evidence supporting the feasibility of cutaneous monitoring of blood glucose level *in vivo* with mouse models. The skin-based glucose sensor provides an accurate and stable glucose monitoring with a wide dynamic range. Because of low immunogenicity of autologous skin transplants, our probe will have superior durability for long term application *in vivo.*

Given the enhanced sensitivity and non-invasive nature, fluorescence-based glucose sensing has been actively explored for continuous glucose monitoring ^29,47^. Measurement of the fluorescence signals causes little or no damage to the host. With future improvement of the probe, such as increased sensor expression in the epidermal cells and using FRET pairs with longer excitation wavelength, we can further reduce the exposure time of skin to fluorescence light source for repeated measurements, thus minimizing the potential photo-damages of the skin. The same platform can be also exploited to develop other biosensors and stably deliver therapeutic proteins *in vivo*, such as GLP1, providing a promising treatment for many human diseases ^13,14^.

Taken together, our study demonstrates that grafted skin stem cells expressing a foreign reporter protein can be efficiently and stably grafted to the host mice with intact immune systems, and can be used to accurately monitor metabolic changes *in vivo*. Skin-based gene therapy has been proposed to treat many diseases, including inherited skin disorders ^19,49-51^ and other terminal or severely disabling diseases ^13-18^. Our study unravels the tempting potential of cutaneous gene therapy for various clinical applications in the future.

## ACKNOWLEDGMENTS

We are very grateful to Dr. Lev Becker at the University of Chicago, Dr. Markus Schober at New York University School of Medicine, and Dr. Elaine Fuchs at the Rockefeller University for sharing reagents and technical assistance. We thank Linda Degenstein at the Transgenic Core Facility, Dr. Lara Leoni at the Animal Imaging Facility, and Drs. Vytas Bindokas and Christine Labno at the Light Microscopy Core Facility of the University of Chicago for excellent technical assistance. The animal studies were carried out in the ALAAC-accredited animal research facility at the University of Chicago. This work was supported by a grant R01-AR063630 from the National Institutes of Health, the Research Scholar Grant (RSG-13-198-01) from the American Cancer Society, and the V scholar award from V foundation to XW.

## STAR Methods

### Contact for Reagent and Resource Sharing

Further information and requests for resources and reagents should be directed to and will be fulfilled by the Lead Contact, Xiaoyang Wu.

### Experimental Model and Subject Details

#### Animals

CD1 mice were obtained from the Transgenic Core Facility at University of Chicago. Athymic nude mice were purchased from Jackson Laboratories. All the mice were housed under pathogen-free conditions in ARC (animal resource center) of University of Chicago under a 12-hour light-dark cycle. The experimental protocol was reviewed and approved by the Institutional Animal Care and Use Committee of University of Chicago.

#### Primary Epidermal Cell Culture

Primary mouse epidermal basal keratinocytes were isolated from the epidermis of newborn mice using trypsin, after prior separation of the epidermis from the dermis by an overnight dispase treatment. Keratinocytes were plated on mitomycin C–treated 3T3 fibroblast feeder cells until passage 5-8. Cells were cultured in E-media supplemented with 15% serum and a final concentration of 0.05 mM Ca^2+^ ^52^.

Primary human epidermal basal keratinocytes were purchased from Invitrogen. Cells were cultured on mitomycin C–treated 3T3 fibroblast feeder cells with E-media supplemented with 15% serum and a final concentration of 0.05 mM Ca^2+^ ^8,9^.

### Method Details

#### Plasmid DNA Constructions

Lentiviral vector encoding *Luciferase* and *H2B-RFP* has been described before ^21,23^. Plasmid encoding *hCas9*-D10A mutant was a gift from George Church, obtained from Addgene (plasmid #41816). Plasmid encoding gRNA expression cassette was constructed with primers: AAG GAA AAA AGC GGC CGC TGT ACA AAA AAG CAG G; and gGA ATT CTA ATG CCA ACT TTG TAC, using gBlock as a template. *Rosa26* –targeting gRNA is constructed with primers: ACA CCG GCA GGC TTA AAG GCT AAC CG, AAA ACG GTT AGC CTT TAA GCC TGC CG, ACA CCG AGG ACA ACG CCC ACA CAC Cg, AAA ACG GTG TGT GGG CGT TGT CCT CG. *AAVS1*–targeting gRNA is constructed with primers: ACA CCG TCA CCA ATC CTG TCC CTA GG, AAA ACC TAG GGA CAG GAT TGG TGA CG, ACA CCG CCC CAC AGT GGG GCC ACT AG, AAA ACT AGT GGC CCC ACT GTG GGG CG. *Rosa26* targeting vector is constructed with pRosa26-GT as template (a gift from Liqun Luo, addgene plasmid 40025) using primers: GAC TAG TGA ATT CGG ATC CTT AAT TAA GGC CTC CGC GCC GGG TTT TGG CG, GAC TAG TCC CGG GGG ATC CAC CGG TCA GGA ACA GGT GGT GGC GGC CC, CGG GAT CCA CCG GTG AGG GCA GAG GAA GCC TTC TAA C, TCC CCC GGG TAC AAA ATC AGA AGG ACA GGG AAG, GGA ATT CAA TAA AAT ATC TTT ATT TTC ATT ACA TC, CCT TAA TTA AGG ATC CAC GCG TGT TTA AAC ACC GGT TTT ACG AGG GTA GGA AGT GGT AC. *AAVS1* targeting vector was constructed with AAVS1 hPGK-PuroR-pA donor (a gift from Rudolf Jaenisch, addgene plasmid 22072) as template using primers: CCC AAG CTT CTC GAG TTG GGG TTG CGC CTT TTC CAA G, CCC AAG CTT CCA TAG AGC CCA CCG CAT CCC C, CAG GGT CTA GAC GCC GGA TCC GGT ACC CTG TGC CTT CTA GTT GC, GGA TCC GGC GTC TAG ACC CTG GGG AGA GAG GTC GGT G, CCG CTC GAG AAT AAA ATA TCT TTA TTT TCA TTA CAT C, GCT CTA GAC CAA GTG ACG ATC ACA GCG ATC. Genotyping primers for CIRPSR mediated knockin: GAG CTG GGA CCA CCT TAT ATT C, GGT GCA TGA CCC GCA AG, GAG AGA TGG CTC CAG GAA ATG.

#### Genome editing and selection of engineered epidermal cells

Human or mouse primary epidermal basal cells were electroporated with a mixture of plasmid DNA containing plasmid encoding *hCas9-*D10A, plasmids encoding the gRNAs targeting *Rosa26* locus for mouse cells or *AAVS1* locus for human cells, and the *GLP1* targeting construct. Electroporation was carried out with BioRad gene pulser using Ingenio electroporation solution. Cells were electroporated using exponential decay mode at 250 Volts and 950 μF. Electroporated cells were immediately suspended in culture medium and plated. Cells were selected with puromycin (2.5μg/ml) 2-3 days post electroporation.

Mouse and human cells were electroporated at passage 8-10. Mouse epidermal cells at this stage can grow in feeder-free condition, whereas human cells are continuously cultured with feeder cells. Limited trypsinization or brief EDTA (20 mM in PBS) treatment was used to remove the feeder cells before suspension of human epidermal cells.

#### Skin organoid culture and transplantation

Decelluralized dermis (circular shape with 1cm diameter) was prepared by EDTA treatment of newborn mouse skin ^24^. 1.5 X 10^6^ cultured keratinocytes were seeded onto the dermis in cell culture insert. After overnight attachment, the skin culture was exposed to air/liquid interface.

For grafting with skin organoids, CD1 males with the ages of 6-8 weeks were anesthetized. A silicone chamber bottom with the interior diameter of 0.8cm and the exterior diameter of 1.5cm was implanted on its shaved dorsal mid-line skin, which was used to hold the skin graft. A chamber cap was installed to seal the chamber right after a piece of graft was implanted. About one week later, the chamber cap was removed to expose the graft to air. A single dose of 0.2mg α-CD4 (GK1.5) and 0.2mg α-CD8 (2.43.1) antibodies was administered intraperitoneally for skin grafting.

#### Cell cycle analysis

Propidium Iodide (PI) staining followed by Flow Cytometry Assay were used to determine the effect of cell cycle profiles. Mouse and human epidermal cells were cultured in two 6cm cell culture dish for 24 hours, respectively. Cells were trypsinized and 1×10^5^ cells from each dish were collected, followed by one PBS wash. Fixation of cells was carried out using 70% (v/v) ice cold ethanol for 1 hour. Then, the fixed cells were centrifuged at 500g at 4°C for 10 minutes, followed by PBS wash for two times. The cells were then treated with 75μg RNAse A in 100μl PBS and incubated at 37°C for 1 hour. After incubation, the cells were collected by centrifuging at 500g at 4°C for 10 minutes, followed by another PBS wash. The cell pellet was re-suspended in 200μl PBS, in addition of PI solution at a final concentration of 25ng/μl. After staining, the cells were analyzed immediately using flow cytometer BD FACSCanto^TM^ II (BD Biosciences, San Jose, CA) with an excitation wavelength at 488 nm and emission at 585 nm. DNA content and histograms of cell cycle distribution were analyzed using FlowJo software, version 10 (FLOWJO LLC, OR).

#### Obesity induced by high fat diet and glucose tolerance test

Male CD-1 mice with skin transplants were housed (5 per cage, ∼8 weeks old) in a central-controlled animal facility for air, humidity and temperature. These mice were fed either a regular chow or an HFD (60% kcal from fats, 20% from carbohydrates, and 20% from proteins) purchased from Bio-Serv (Frenchtown, NJ). Body weight and food intake were measure biweekly.

For Glucose tolerance tolerance test, an intraperitoneal glucose tolerance test (IPGTT) was performed on mice fed an HFD for 10 weeks. Mice were fasted for 6 h before the test. Animals were injected (1 g/kg glucose/body weight, i.p.) with glucose dissolved in saline, and blood glucose was measured at 0, 10, 20, 30, 60 and 90 minutes using glucose test strips and glucose meters.

To induce hypoglycemia, CD1 mice with skin grafts were fasted for 4 h and injected (2 U/kg, i.p.) with insulin purchased from Sigma (St. Louis, MO). Blood glucose levels were determined thereafter at 0, 15, 30, 45, and 60 minutes.

To examine feeding-caused glucose level change, animals were fasted overnight, and then fed with regular chow diet for 30 minutes. Glucose level was measured at 0, 30, 60, and 120 minutes with glucose test strips and fluorescence changes in the skin grafts. GLP1 level was measured by ELISA at 0-6 hours post feeding.

#### Intravital imaging of mice

Optical imaging was performed in the integrated small animal imaging research resource (iSAIRR) at the University of Chicago. Bioluminescence images were acquired on an IVIS Spectrum (Caliper Life Sciences, Alameda, CA) after animal was injected with luciferin (100mg/kg). Acquisition and image analysis were performed with Living Image 4.3.1 software.

We used the ImageJ PixFRET plugin for image analysis and calculation of FRET ratio. The parameters in the “Donor Model” and “Acceptor Model” were calculated based on the donor and acceptor single control images. The following parameters were used for the image processing:

1. Donor model: Background Donor:FRET=2.28, Donor=14.35. Model Donor:Gaussian Blur=2.0; Linear: a=0.18964, b=-0.02746.
2. Acceptor model: Background Donor:FRET=1.21, Acceptor=3.93. Model Donor:Gaussian Blur=2.0; Linear: a=0.26711, b=-0.07069
3. FRET model: Gaussian blue=0, Threshold=1.5, Output=FRET/Donor.

Mean value of FRET efficiency was calculated from the output images using the Analyze/Measure function of Image J.

#### Histology and Immunofluorescence

Skin or wound samples were embedded in OCT, frozen, sectioned, and fixed in 4% formaldehyde. For paraffin sections, samples were incubated in 4% formaldehyde at 4°C overnight, dehydrated with a series of increasing concentrations of ethanol and xylene, and then embedded in paraffin. Paraffin sections were rehydrated in decreasing concentrations of ethanol and subjected to antigen unmasking in 10 mM Citrate, pH 6.0. Sections were subjected to hematoxylin and eosin staining or immunofluorescence staining as described ^53^. Antibodies were diluted according to manufacturer’s instruction, unless indicated.

### Quantification and Statistical Analysis

Statistical analysis was performed using Excel or OriginLab software. Box plots are used to describe the entire population without assumptions on the statistical distribution. A student *t* test was used to assess the statistical significance (P value) of differences between two experimental conditions (2 tailed distribution unless specified). All experiments were repeated at least three times, unless otherwise specified. For all figures, statistical tests are justified and meet the assumption of the tests. The variance between different test groups that are being statistically compared is similar.

For all the experiments, the sample size was chosen based upon our preliminary test and previous research. There is no sample exclusion for all the *in vitro* analysis. For *in vivo* experiments, animals that died before the end of the experiment were excluded. The exclusion criterion is pre-established. No randomization or blinding was used in this study.

### Data and Software Availability

A complete list of software for data analysis and processing can be found in the Key Resource Table.

## References

1 Kim, D. H., Ghaffari, R., Lu,N. & Rogers, J. A. Flexible and stretchable electronics for biointegrated devices. Annual review of biomedical engineering 14, 113–128, doi:10.1146/annurev-bioeng-071811-150018 (2012).

2 Choi, S., Lee, H., Ghaffari, R., Hyeon, T. & Kim, D. H. Recent Advances in Flexible and Stretchable Bio-Electronic Devices Integrated with Nanomaterials. Advanced materials 28, 4203–4218, doi:10.1002/adma.201504150 (2016).

3 Hsu, P. D., Lander, E. S. & Zhang, F. Development and applications of CRISPR-Cas9 for genome engineering. Cell 157, 1262–1278, doi:10.1016/j.cell.2014.05.010 (2014).

4 Maeder, M. L. & Gersbach, C. A. Genome-editing Technologies for Gene and Cell Therapy. Mol Ther 24, 430–446, doi:10.1038/mt.2016.10 (2016).

5 Wright, A. V., Nunez, J. K. & Doudna, J. A. Biology and Applications of CRISPR Systems: Harnessing Nature's Toolbox for Genome Engineering. Cell 164, 29–44, doi:10.1016/j.cell.2015.12.035 (2016).

6 Blanpain, C. & Fuchs, E. Epidermal stem cells of the skin. Annu Rev Cell Dev Biol 22, 339– 373, doi:10.1146/annurev.cellbio.22.010305.104357 (2006).

7 Watt, F. M. Mammalian skin cell biology: at the interface between laboratory and clinic. Science 346, 937–940, doi:10.1126/science.1253734 (2014).

8 Rasmussen, C., Thomas-Virnig,C. & Allen-Hoffmann, B. L. Classical human epidermal keratinocyte cell culture. Methods Mol Biol 945, 161–175, doi:10.1007/978-1-62703-125-7_11 (2013).

9 Rheinwald, J. G. & Green, H. Serial cultivation of strains of human epidermal keratinocytes: the formation of keratinizing colonies from single cells. Cell 6, 331–343 (1975).

10 Rheinwald, J. G. & Green, H. Epidermal growth factor and the multiplication of cultured human epidermal keratinocytes. Nature 265, 421–424 (1977).

11 Carsin, H. et al. Cultured epithelial autografts in extensive burn coverage of severely traumatized patients: a five year single-center experience with 30 patients. Burns: journal of the International Society for Burn Injuries 26, 379–387 (2000).

12 Coleman, J. J., 3rd & Siwy, B. K. Cultured epidermal autografts: a life-saving and skin-saving technique in children. Journal of pediatric surgery 27, 1029–1032 (1992).

13 Christensen, R., Jensen, U. B. & Jensen, T. G. Skin genetically engineered as a bioreactor or a 'metabolic sink'. Cells, tissues, organs 172, 96–104, doi:65612 (2002).

14 Del Rio, M., Gache, Y., Jorcano, J. L., Meneguzzi, G. & Larcher, F. Current approaches and perspectives in human keratinocyte-based gene therapies. Gene therapy 11 Suppl 1, S57–63, doi:10.1038/sj.gt.3302370 (2004).

15 Fakharzadeh, S. S., Zhang, Y., Sarkar, R. & Kazazian, H. H., Jr. Correction of the coagulation defect in hemophilia A mice through factor VIII expression in skin. Blood 95, 2799–2805 (2000).

16 Fenjves, E. S., Gordon, D. A., Pershing, L. K., Williams, D. L. & Taichman, L. B. Systemic distribution of apolipoprotein E secreted by grafts of epidermal keratinocytes: implications for epidermal function and gene therapy. Proc Natl Acad Sci U S A 86, 8803–8807 (1989).

17 Gerrard, A. J., Hudson, D. L., Brownlee, G. G. & Watt, F. M. Towards gene therapy for haemophilia B using primary human keratinocytes. Nature genetics 3, 180–183, doi:10.1038/ng0293-180 (1993).

18 Morgan, J. R., Barrandon, Y., Green, H. & Mulligan, R. C. Expression of an exogenous growth hormone gene by transplantable human epidermal cells. Science 237, 1476–1479 (1987).

19 Sebastiano, V. et al. Human COL7A1-corrected induced pluripotent stem cells for the treatment of recessive dystrophic epidermolysis bullosa. Science translational medicine 6, 264ra163, doi:10.1126/scitranslmed.3009540 (2014).

20 Collins, M. & Thrasher, A. Gene therapy: progress and predictions. Proceedings. Biological sciences / The Royal Society 282, doi:10.1098/rspb.2014.3003 (2015).

21 Liu, H. et al. Regulation of Focal Adhesion Dynamics and Cell Motility by the EB2 and Hax1 Protein Complex. J Biol Chem 290, 30771–30782, doi:10.1074/jbc.M115.671743 (2015).

22 Yue, J., Gou, X., Li, Y., Wicksteed, B. & Wu, X. Engineered Epidermal Progenitor Cells Can Correct Diet-Induced Obesity and Diabetes. Cell Stem Cell 21, 256–263 e254, doi:10.1016/j.stem.2017.06.016 (2017).

23 Yue, J. et al. In vivo epidermal migration requires focal adhesion targeting of ACF7. Nat Commun 7, 11692, doi:10.1038/ncomms11692 (2016).

24 Prunieras, M., Regnier, M. & Woodley, D. Methods for cultivation of keratinocytes with an air-liquid interface. J Invest Dermatol 81, 28s–33s (1983).

25 Ahima, R. S. Digging deeper into obesity. J Clin Invest 121, 2076–2079, doi:10.1172/JCI58719 (2011).

26 Oliver, N. S., Toumazou, C., Cass, A. E. & Johnston, D. G. Glucose sensors: a review of current and emerging technology. Diabetic medicine: a journal of the British Diabetic Association 26, 197–210, doi:10.1111/j.1464-5491.2008.02642.x (2009).

27 Pickup, J. Developing glucose sensors for in vivo use. Trends in biotechnology 11, 285–291, doi:10.1016/0167-7799(93)90016-3 (1993).

28 Wang, J. Electrochemical glucose biosensors. Chemical reviews 108, 814–825, doi:10.1021/cr068123a (2008).

29 Pickup, J. C., Hussain, F., Evans, N. D., Rolinski, O. J. & Birch, D. J. Fluorescence-based glucose sensors. Biosensors & bioelectronics 20, 2555–2565, doi:10.1016/j.bios.2004.10.002 (2005).

30 Jeffery, C. J. Engineering periplasmic ligand binding proteins as glucose nanosensors. Nano reviews 2, doi:10.3402/nano.v2i0.5743 (2011).

31 Fehr, M., Lalonde, S., Lager, I., Wolff, M. W. & Frommer, W. B. In vivo imaging of the dynamics of glucose uptake in the cytosol of COS-7 cells by fluorescent nanosensors. J Biol Chem 278, 19127–19133, doi:10.1074/jbc.M301333200 (2003).

32 Saxl, T., Khan, F., Ferla, M., Birch, D. & Pickup, J. A fluorescence lifetime-based fibre-optic glucose sensor using glucose/galactose-binding protein. The Analyst 136, 968–972, doi:10.1039/c0an00430h (2011).

33 Teasley Hamorsky, K., Ensor, C. M., Wei, Y. & Daunert, S. A bioluminescent molecular switch for glucose. Angewandte Chemie 47, 3718–3721, doi:10.1002/anie.200704440 (2008).

34 Tian, Y. et al. Structure-based design of robust glucose biosensors using a Thermotoga maritima periplasmic glucose-binding protein. Protein science: a publication of the Protein Society 16, 2240–2250, doi:10.1110/ps.072969407 (2007).

35 Veetil, J. V., Jin, S. & Ye, K. A glucose sensor protein for continuous glucose monitoring. Biosensors & bioelectronics 26, 1650–1655, doi:10.1016/j.bios.2010.08.052 (2010).

36 Amiss, T. J., Sherman, D. B., Nycz, C. M., Andaluz, S. A. & Pitner, J. B. Engineering and rapid selection of a low-affinity glucose/galactose-binding protein for a glucose biosensor. Protein science: a publication of the Protein Society 16, 2350–2359, doi:10.1110/ps.073119507 (2007).

37 Kotterman, M. A., Chalberg, T. W. & Schaffer, D. V. Viral Vectors for Gene Therapy: Translational and Clinical Outlook. Annual review of biomedical engineering 17, 63–89, doi:10.1146/annurev-bioeng-071813-104938 (2015).

38 Kustikova, O., Brugman, M. & Baum, C. The genomic risk of somatic gene therapy. Seminars in cancer biology 20, 269–278, doi:10.1016/j.semcancer.2010.06.003 (2010).

39 Cox, D. B., Platt, R. J. & Zhang, F. Therapeutic genome editing: prospects and challenges. Nat Med 21, 121–131, doi:10.1038/nm.3793 (2015).

40 Hotta, A. & Yamanaka, S. From Genomics to Gene Therapy: Induced Pluripotent Stem Cells Meet Genome Editing. Annual review of genetics 49, 47–70, doi:10.1146/annurev-genet-112414-054926 (2015).

41 Ran, F. A. et al. Double nicking by RNA-guided CRISPR Cas9 for enhanced genome editing specificity. Cell 154, 1380–1389, doi:10.1016/j.cell.2013.08.021 (2013).

42 Schober, M. & Fuchs, E. Tumor-initiating stem cells of squamous cell carcinomas and their control by TGF-beta and integrin/focal adhesion kinase (FAK) signaling. Proc Natl Acad Sci U S A 108, 10544–10549, doi:1107807108 [pii]10.1073/pnas.1107807108 (2011).

43 Sandoval, D. A. & D'Alessio, D. A. Physiology of proglucagon peptides: role of glucagon and GLP-1 in health and disease. Physiological reviews 95, 513–548, doi:10.1152/physrev.00013.2014 (2015).

44 Kumar, M., Hunag, Y., Glinka, Y., Prud'homme, G. J. & Wang, Q. Gene therapy of diabetes using a novel GLP-1/IgG1-Fc fusion construct normalizes glucose levels in db/db mice. Gene therapy 14, 162–172, doi:10.1038/sj.gt.3302836 (2007).

45 Steiner, M. S., Duerkop, A. & Wolfbeis, O. S. Optical methods for sensing glucose. Chemical Society reviews 40, 4805–4839, doi:10.1039/c1cs15063d (2011).

46 Tian, K., Prestgard, M. & Tiwari, A. A review of recent advances in nonenzymatic glucose sensors. Materials science & engineering. C, Materials for biological applications 41, 100–118, doi:10.1016/j.msec.2014.04.013 (2014).

47 Vaddiraju, S., Burgess, D. J., Tomazos, I., Jain, F. C. & Papadimitrakopoulos, F. Technologies for continuous glucose monitoring: current problems and future promises. Journal of diabetes science and technology 4, 1540–1562 (2010).

48 Heo, Y. J. & Takeuchi, S. Towards smart tattoos: implantable biosensors for continuous glucose monitoring. Advanced healthcare materials 2, 43–56, doi:10.1002/adhm.201200167 (2013).

49 Chen, M. et al. Restoration of type VII collagen expression and function in dystrophic epidermolysis bullosa. Nature genetics 32, 670–675, doi:10.1038/ng1041 (2002).

50 Ortiz-Urda, S. et al. PhiC31 integrase-mediated nonviral genetic correction of junctional epidermolysis bullosa. Human gene therapy 14, 923–928, doi:10.1089/104303403765701204 (2003).

51 Ortiz-Urda, S. et al. Stable nonviral genetic correction of inherited human skin disease. Nat Med 8, 1166–1170, doi:10.1038/nm766 (2002).

52 Wu, X., Kodama,A. & Fuchs, E. ACF7 regulates cytoskeletal-focal adhesion dynamics and migration and has ATPase activity. Cell 135, 137–148, doi:S0092-8674(08)01011-8 [pii]10.1016/j.cell.2008.07.045 (2008).

53 Guasch, G. et al. Loss of TGFbeta signaling destabilizes homeostasis and promotes squamous cell carcinomas in stratified epithelia. Cancer Cell 12, 313–327, doi:S1535- 6108(07)00239-5 [pii] 10.1016/j.ccr.2007.08.020 (2007).

